# Increased spontaneous Ca^2+^ activity in Cardiac Purkinje cells after myocardial infarction; A consequence of a dramatic shift of SERCA isoforms as potential adaptation to acute ischemia?

**DOI:** 10.1101/2023.10.23.563678

**Authors:** Bruno D. Stuyvers, Penelope A. Boyden, Ruhul Amin, Louisa Wiede, Jules Doré, Henk E.D.J. ter Keurs, Wen Dun, Yunbo Guo, Michel Haissaguerre, Meleze Hocini, Fabien Brette, Olivier Bernus, Sebastien Chaigne, Virginie Loyer, Jerome Naulin, Bruno Quesson, Akinola Alafiatayo, Touati Benoukraf

## Abstract

**Background:** Studies of Purkinje cells (Pcells) from canine hearts have suggested an increase of Ca^2+^-release by the sarcoplasmic reticulum (SR) but also reported a potential augmentation of SR-Ca^2+^-uptake after MI. Abnormal increase of SR-Ca^2+^-uptake in heart cells is novel and contrasts with the reduction of this function in cells of failing heart. Our study examined the origin of this increased SR-Ca^2+^-uptake by considering a change in SR-Ca^2+^ pump (SERCA2) expression in Purkinje fibers (PFs) post MI.

**Methods:** Pcells were isolated from canine hearts 48Hrs post MI. Intracellular Ca^2+^-activity was captured by confocal microscopy. Purkinje-typical Ca^2+^ events were analyzed to probe the regional Ca^2+^-dynamics within Pcells. A Purkinje-specific numerical model assisted in the interpretation of Ca^2+^-anomalies detected in Pcells Ca^2+^-transients. SR-Ca^2+^-uptake system was studied by immunofluorescence in Pcells from canine, ovine and human hearts post MI. SERCA protein and gene expressions in PFs and myocardium were measured by Western Blots and RT-qPCR in a classical porcine model of MI.

**Results:** 48Hrs after MI, Pcells showed 60% increase in spark-rate and 37% acceleration of Ca^2+^ wave decay. In the model of normal wave, 35% increase of Ca^2+^-uptake rate reproduced the actual post-MI wave alterations. In apparent contrast with increased Ca^2+^-uptake rate, SERCA2 protein expression was reduced in canine, sheep, and human Pcells after MI. In pig MI model, the protein level of cardiac-specific SERCA2-splicing variant SERCA2a was reduced by 52% in the whole infarcted ventricle whereas the “non-cardiac” SERCA2b level was increased by 120%. In the infarcted regions, PFs showed 30% downregulation of SERCA2a gene expression and 630% upregulation of SERCA2b.

**Conclusion:** Our results confirm that elevated spontaneous Ca^2+^-activity in post-MI PFs is due to increased SR-Ca^2+^-uptake within Pcells. Data suggest that a replacement of “cardiac” SERCA2a by the “non-cardiac” SERCA2b sub-isoform in cardiac cells in response to ischemia is implicated in this alteration.

## Introduction

Experimental and clinical evidence attributes the initiation of ischemic tachyarrhythmias to non-driven depolarizations of Purkinje fibers ^1–8^. During the early phase post myocardial infarction (MI), the cells of subendocardial Purkinje fibers (Pcells) survive in the infarct ^9,10^, but exhibit afterdepolarizations (DADs) and non-driven action potentials (APs) that can initiate ventricular tachycardias ^3,9^. Pro-arrhythmic depolarizations have been attributed to abnormal intracellular Ca^2+^ handling ^3,4,11–13^. In heart cells including Pcells, large Ca^2+^ waves can activate depolarizing currents via Na-Ca exchange (NCX) and potentially trigger full action potentials ^3,11,14–17^. In Pcells, the probability of a large Ca^2+^ wave increases with the incidence of smaller spontaneous Ca^2+^ events like sparks and wavelets ^18^. An augmentation of such small Ca^2+^ events was found in canine Pcells within two days following the ligature of the LAD coronary artery, thus supporting the causal relationship between increased spontaneous Ca^2+^ activity (SCA) in Pcells and Purkinje arrhythmicity in post MI hearts ^19,20^. The process whereby myocardial ischemia potentiates spontaneous Ca^2+^-releases in Pcells remains unknown.

Ca^2+^ content of the sarcoplasmic reticulum (SR) is a major determinant of spontaneous release of Ca^2+^ in cardiac cells ^21^. We have reported that SR-Ca^2+^ content of canine Pcells remains unchanged 48Hrs after MI ^20^. Thus SR-Ca^2+^ overload is not involved in the increased spontaneous Ca^2+^-release. These findings rather suggest an involvement of altered SR-Ca^2+^ trans-membrane transport systems. We have previously reported that the curve rate of spontaneous Ca^2+^ events versus Ca^2+^ concentration (pCa) in Pcells was shifted upward after MI ^20^. This finding may indicate an increase of Ca^2+^ sensitivity of SR-Ca^2+^-release channels. Nevertheless, in the same study, a faster Ca^2+^ decay in the “micro-Ca^2+^ transients” was observed in MI Pcells. This observation is consistent with faster cytosolic Ca^2+^ removal by the Ca^2+^ extrusion systems. The absence of transverse (T-)tubules in Pcells of large mammalian species ^22,23^ implies a modest contribution of sarcolemmal Ca^2+^-ATPases and NCX to cytosolic Ca^2+^ extrusion in the core of Pcells. Furthermore, it predicts that SR-Ca^2+^ pumps dominate the control of basal Ca^2+^ level in this region. Thus it is reasonable to conclude that faster cytosolic Ca^2+^ removal reported in ^20^ reflected an increased activity of SR-Ca^2+^-ATPases (SERCA) in MI Pcells. Therefore, we hypothesized that an alteration of SERCA activity takes place in Pcells shortly after the onset of myocardial ischemia and, as such, contributes to the elevation of pro-arrhythmic SCA in Purkinje fibers during the early phase of the infarction.

SR-Ca^2+^-release function was assessed in Pcells of canine heart 48Hrs post MI through the analysis of typical Ca^2+^ transients previously identified in Pcells ^18,22^. The SR-Ca^2+^-uptake system was examined by determining the density distribution of SERCA pumps and regulatory protein phospholamban (PLB). To evaluate the model dependence and translational relevance of our results from canine hearts, experiments were implemented or reproduced in sheep, human and pig hearts.

Our study confirms that the SR-Ca^2+^-uptake in Pcells shows a marked increase within two days from the onset of ischemia. In these same cells, we report here a dramatic re-arrangement of SERCA pumps. In addition, SERCA pump expression in Purkinje fibers from the infarcted ventricles revealed a large upregulation of non-cardiac SERCA2b isoform. The relationship between increased SR-Ca^2+^-uptake in Pcells and alteration of SERCA expression in ischemic heart is discussed.

## Methods

### Preparation of Purkinje cells and models of Myocardial Infarctions

Acute myocardial ischemia was induced in adult mongrel dog by permanent double ligation technique of left anterior descending (LAD) coronary artery as in ^9,24^. The hearts were explanted and Pcells were isolated 48Hrs after the ligation. Pcells preparation was modified from original methods of Sheets *et al.* ^25^ and Boyden *et al.* ^9^. Briefly, sub-endocardial Purkinje strands were dissected from the left ventricular chamber. After incubation of the strands with collagenase (Type II, Worthington) at 37°C, pH 6.7 for 30-40 min, the cells were mechanically dispersed with the same medium with no collagenase, then filtered and finally re-suspended in HEPES-buffered medium with 2mM Ca^2+^ and pH 7.3. The cells were used the same day for intracellular Ca^2+^ imaging and/or protein immuno-staining. Two different groups of cells, normal (control) and “MI” Pcells, were prepared respectively from Purkinje fibers of non-infarcted hearts and hearts with 48hrs MI. MI Pcells were isolated from Purkinje fibers located in the subendocardial region of the infarct and border zone (BZ).

We compared our results from the canine model with those from an ovine (sheep) model where MI was induced by angiographic method as described in **Fig.5A**.

Pcells have also been collected from human hearts not suitable for transplant. Normal hearts, i.e., with no sign of anatomical or metabolic abnormalities and excluded from transplant program based only on incompatibility of patient profile, were compared with one heart with recent typical ischemic MI (see supplement **Fig.6S**). Collection and handling of human hearts followed human ethic policies associated with *CADENCE* project involving the *groupe hospitalier Pellegrin-CHU de Bordeaux*, LIRYC Institute, and University of Bordeaux, France.

Pcells from canine, ovine, and human hearts were prepared using identical procedure. *Utilization of animals as described in the protocols has been approved by institutional animal care committees of Memorial University of Newfoundland, University of Calgary, Columbia University and Université de Bordeaux, and complied with policies of experimental animal practice of Canada, USA, and France*.

### Study of intracellular Ca^2+^ dynamics

Freshly isolated P-cells were placed in a field-stimulated experimental chamber and continuously perfused with 2mM Ca^2+^ standard HEPES solution (Tyrode) at 37°C (pH 7.3) for 5 min. The perfusion was interrupted, and coverslip-adherent cells were incubated with 5 µM Fluo4-AM in the same solution for 20-30 min. Then, the cells were rinsed and left in the dark for 10 min. The cells were field stimulated at low frequency (0.1Hz) to uniformly activate the intracellular Ca^2+^ cycling throughout the cell preparation.

A Fluo-4-loaded Pcell was selected and illuminated at 388 nm. Emitted fluorescence was simultaneously collected at 586 nm. Ca^2+^ dependent fluorescence of Fluo-4 was captured by line scanning confocal microscopy (LSCM) at 333 scans/sec or 2-dimensional spinning disk confocal microscopy (2DCM) at 30 frames per second (fps) as described elsewhere ^18,20,22,26^. The relative free-Ca^2+^ concentration ([Ca^2+^]) was given by the pixel-to-pixel ratio F/Fo, where F and Fo were the instantaneous fluorescence intensity and basal fluorescence intensity, respectively. As measured previously under identical experimental conditions, F/Fo = 1 was equivalent to [Ca^2+^] = 100 nml.L^-1 18^. [Ca^2+^] variations were given by the ratio F-Fo/Fo (“ΔF/Fo”). Image processing and Ca^2+^ transient analyses were carried out using custom IDL programs (L3Harris Geospatial Solutions, Inc. USA) and modules of the open-source NIH software ImageJ (see supplement Figs.1S).

### Comparative modeling of Ca^2+^-transients

To assist in the interpretation of data from Pcells, we used our previous model based on the combination of Ca^2+^-release, diffusion, binding and re-uptake in a 2D-array of “Ca^2+^ nodes” ^18,27^. The model’s suitability for our current study was tested by reproducing (1) the individual Ca^2+^ transients measured by LSCM (see supplement **Fig.3S**) and (2) the 2 averaged Ca^2+^ transients of normal and MI Pcells, respectively (see **Fig. 2A**). The accuracy of the simulations was evaluated by linear regression between computational and experimental data. The initial conditions of the model were described previously ^27^; see also “overview of the model” in supplement.

### Immuno-characterization of SR-Ca^2+^-uptake system in single Pcells

The protein expressions of SERCA pump and phospholamban (PLB) were studied in situ by immunofluorescence and confocal microscopy in single Pcells and intact Purkinje fibers from canine normal (non-infarcted) ventricles and ventricles post 48Hrs ischemic MI. Single Pcells were isolated within 1Hr upon heart explant. They were extensively washed with PBS, then fixed with 4% formaldehyde and permeabilized with 0.3% Triton X100, and finally incubated overnight at 4°C with the primary antibodies mouse anti-SERCA2 (monoclonal 1:500, Affinity BioReagents), and rabbit anti-pPLBSer16 (1:1000, Badrilla). The next day, the cells were incubated with the secondary fluorescent antibodies Alexa Fluor® 488 Goat Anti-Rabbit IgG (H+L) and Alexa Fluor® 568 Goat Anti-Mouse IgG (H+L) (Life Technologies). Ovine (**Fig. 5**) and human Pcells (supplement **Fig.5S**) were treated with monoclonal mouse SERCA2 antibody (clone 2A7-A1; Thermo Fisher; see supplement) and secondary fluorescent antibody Alexa Fluor® 568 Goat Anti-Mouse IgG (H+L) (Life Technology) following the same procedure. Immunofluorescence-labeled signals were sampled with 488 nm and 594 nm excitation, respectively, using a Zeiss 510 META LSCM (canine Pcells) or an Olympus Fluoview 1000 LSCM (ovine and human Pcells). Both LSCMs were equipped with 60x Fluo/1.4 oil immersion objective.

### Immuno-characterization of SR-Ca^2+^-uptake system in Purkinje fibers

Purkinje fibers were dissected immediately after heart explant and embedded in O.C.T. (Tissue-Tek). Cryostat-sectioned (12µm) slices were used for immunofluorescence as described in ^28^. Primary SERCA2 and pPLBSer16 antibodies were identical to those used for canine single cells. Sections of Purkinje fibers were examined using a Zeiss 510 NLO multiphoton confocal microscope with 40x Fluor/1.3 oil objective. For quantification, images were converted to black and white using the *threshold* function and *calculate black-to-white* Macro in Image J. The number of white pixels was determined per slice. The amount of labeled protein was expressed as the ratio of white pixels per total cell area (determined through standardized thresholding). Co-localization of signals was determined using the *RGB Co-localization* macro in ImageJ. Results were expressed as pPLB/SERCA2a ratio.

### Quantification of SERCA2a/b transcripts: real-time quantitative reverse transcription–polymerase chain reaction

Total RNA was isolated from Purkinje strands (**Fig. 7A**) using PureLink^TM^ RNA Mini kit (Ref# 12183018A, ThermoFisher Scientific, USA) and quantified with an Agilent, 2100 Bioanalyzer (Agilent Technologies, Lithuania) and RNA Nano Chip (Agilent Technologies, USA). The RNA was reverse transcribed into cDNA with High-Capacity cDNA Reverse Transcription Kit (Ref# 4368814, ThermoFisher Scientific, Lithuania). The qPCR amplification reaction was performed in a 7500 Fast Real-Time PCR system from Applied Biosystems. A 20 µl reaction mixture contained Power SYBR® Green PCR master mix (Ref# 4367659, Thermo Fisher Scientific, UK), 200 nM custom primers (see supplement), and cDNA (1ng RNA equivalent). All samples were quantified in triplicate to ensure the reproducibility and reliability of single values, and normalized to the housekeeping gene, GAPDH (Ss03373286_u1). Relative mRNA levels of SERCA2a and SERCA2b in Purkinje fibers from normal and infarcted ventricles were quantified using the 2^-ΔCT^ method ^29^. In MI hearts, transcript levels were measured in Purkinje fibers from total infarcted LV and, specifically, from infarct region and border zone (BZ). (See supplement for complete method description).

### Ventricular SERCA protein analysis: Western Blots

Since the amount of accessible Purkinje strands in the ventricle is notoriously low, comparison of SERCA2a and SERCA2b protein levels in Purkinje fibers between infarcted and normal ventricles was not possible by immunoblot. However, SERCA2a/2b protein level was evaluated in samples of myocardial wall (including trans-mural Purkinje tissue) of normal and infarcted ventricles.

Myocardial tissues were collected from ventricles immediately after heart extraction. Tissue samples (∼100mg) were flash frozen in liquid nitrogen and stored at -80°C until processed.

Equal amounts of proteins were separated on 8% reducing SDS-PAGE and transferred to Nitrocellulose membranes (Ref# 162-0112, BIO-RAD, Germany). Primary antibodies anti-SERCA2a and anti-SERCA2b targeted the specific ends of SERCA2a and SERCA2b (see supplement). Each antibody specificity was tested separately on porcine tissues by comparing blots from (SERCA2a dominant) myocardium with blots from (SERCA2b dominant) lung tissue (see supplement **Fig.7S**). Membranes were washed extensively prior to incubation with goat anti-rabbit IgG (Ref# HAF008, R&D SYSTEMS, USA) HRP conjugated secondary antibody. Relative protein levels of SERCA2a and SERCA2b were normalized to GAPDH levels for each sample based on the lower molecular mass portion of the same blot probed for GAPDH. (See supplement for complete method description)

### Statistics

Selection or rejection of cells was dictated by the clarity of the fluorescence signals, apparent cell integrity and, in live Pcells, cell response to electric stimulation. Time course parameters of Ca^2+^ transients were averaged for normal and MI groups of cells. Results were expressed as mean ± SEM and compared with unpaired t-test and multi-parametric variance analysis (ANOVA). In RT-PCR and Western Blot analyses, Brown-Forsythe and Welsh ANOVA tests were used for multiple comparisons of data with Gaussian distribution. Non-parametric Kruskal-Wallis test was used for data with non-Gaussian distribution (SERCA2b transcripts). In the modeling section, the degree of correlation between experimental and simulated data was evaluated by linear regression.

## Results

### Alterations of spontaneous Ca^2+^ transients in Pcells after MI

Spontaneous Ca^2+^ activity (SCA) was captured in Fluo-4 loaded Pcells by fluorescence confocal microscopy. The basal fluorescence intensity was steady, and no sign of photo-oxidative alteration was detected during Ca^2+^ imaging sequences. Under our experimental conditions (2mM Ca^2+^, 35°C, pH 7.4), SCA was visible in the form of Ca^2+^ sparks and Ca^2+^ waves in unstimulated condition and between stimulations in Pcells of normal and infarcted ventricles. We compared the intensity of spontaneous SR-Ca^2+^-release between normal and “MI” Pcells by measuring the spark firing rate of individual Ca^2+^-release sites identified along (randomly positioned) 25-50 µm scanlines (**Fig.1A**.a). Consistent with the increase of overall spark rate reported previously ^20^, we found that, on average, Ca^2+^-release sites fired sparks at 60% larger frequency in MI Pcells compared to normal Pcells (**Fig.1A**.b). No difference in the density of active sites was detected between the 2 groups (**Fig.1A**.c).

**FIGURE 1.**
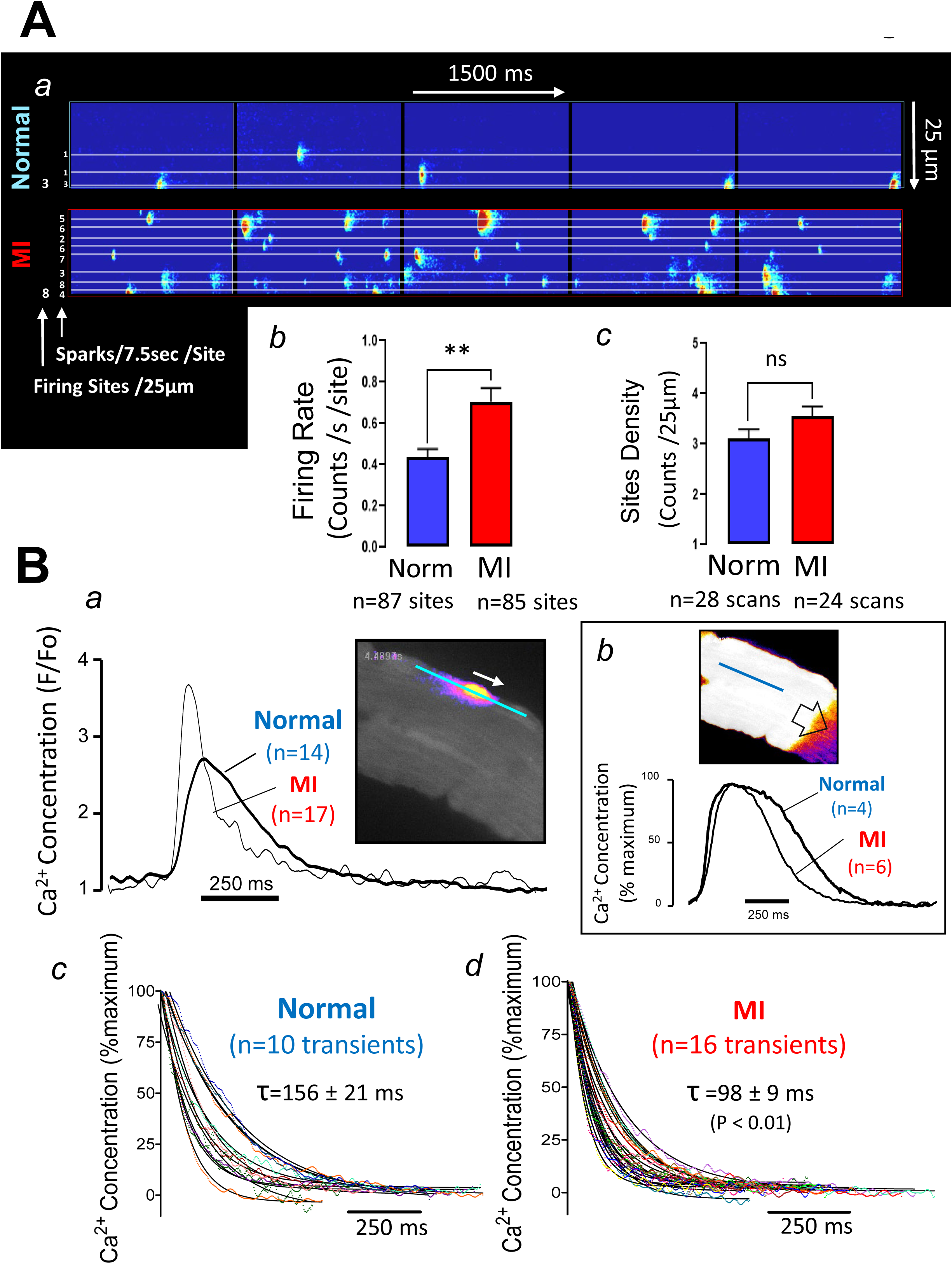
Comparison of spontaneous Ca^2+^ events in dog Pcells from normal heart and heart with 48Hrs MI. **A: Spark activity captured by LSCM in normal and MI Pcells:** panel a illustrates representative Ca^2+^ spark activity along 25 µm line-scan during 1.5s in normal and MI Pcells; panel b represents the number of sites firing sparks (left) and the firing frequency at each site (right); both parameters were measured as indicated in panel a and were compared between normal and MI Pcells in panels b and c, respectively. **B: Averaged time course analysis in dog normal and MI Pcells.** Time courses of 14 normal (6 cells) and 17 MI (6 cells) wavelets were measured as indicated in Figs.1SA (supplement) and averaged in panel a; inset: representative example of one wavelet captured by 2DCM and measured by LSCM (blue line). Averages of CWW time course in normal and MI Pcells were compared in panel b; inset: example of CWW (captured in the same cell shown in inset of panel a). Panels c and d represent the decay time course of 10 and 16 wavelets captured respectively from 6 normal (c) and 7 MI (d) Pcells; a model of exponential decline was used to fit each individual curve and calculate the time constant (τ) of the decay. Individual time constants were averaged and compared as indicated on the graphs of panel c.

Typical cell wide Ca^2+^ waves (CWWs; **Fig.1B**.b) were observed at low rate (approximately one wave every 4-6 seconds) in both normal and MI Pcells. Our wave time course analysis (see supplement **Fig.1S**) revealed a significant decrease of CWW duration with no modification of wave amplitude after MI. This wave “shortening” was due to a 24% reduction of the transient decay (**Fig.1B**.b; Tab. 1) and the typical plateau of CWWs reported in normal Pcells ^18^ was absent in the majority of selected MI Pcells (see supplement **Fig.2S**). In contrast to the large CWWs, Ca^2+^ wavelets are small and localized Ca^2+^ transients (**Fig.1B**.a) propagating over short distances in peripheral regions of Pcells ^18,22^. In MI Pcells, the decay duration of wavelets was significantly reduced by 25% while their amplitude was 50% larger compared to normal Pcells (**Fig.1B**.a, Tab. 1). The time constant (τ) of the decay was decreased by 37% (**Fig.1B**.c,d; Tab. 1) indicating a marked acceleration of cytosolic Ca^2+^ removal in the cell periphery.

**Table 1.** Principal characteristics of Ca^2+^ waves in dog normal and MI Purkinje cells. Time course parameters of wavelets and cell wide waves (CWWs) were measured as explained in FIG.1S (see supplement), from line-scans randomly positioned in the cells; results are expressed as mean ± SEM and n represent the number of transients; unpaired t-test was used to compare values measured from waves captured from normal and MI Pcells, respectively; significant difference was considered when P<0.01. Maximal Amp: maximal amplitude of the transient (measured in Fluo4 F/Fo ratio unit); Duration: duration of the transient; Trise: time to rise to maximum; Tdecay: transient decay duration; τ: time constant of the decay.

| Wavelets | Maximal amp<br>*F/Fo* | Duration<br>*ms* | Trise<br>*ms* | Tdecay<br>*ms* | $\tau$<br>*ms* |
| --- | --- | --- | --- | --- | --- |
| **Normal**<br>(6 hearts)<br>(mean ± SE) | 2.5 ± 0.1<br>(n=83 transients) | 154 ± 8<br>(n=83) | 48 ± 3<br>(n=83) | 106 ± 6<br>(n=83) | 156 ± 21<br>(n=10) |
| **MI**<br>(7 hearts)<br>(mean ± SE) | 3.3 ± 0.1*<br>(n=126 transients) | 118 ± 4*<br>(n=126) | 38 ± 2*<br>(n=126) | 80 ± 3*<br>(n=126) | 98 ± 9 *<br>(n=16) |
| **MI vs Normal**<br>*Difference*<br>(%Normal) | *P*<0.0001<br>0.8 ± 0.2<br>(32%) | *P*<0.0001<br>-36 ± 8<br>(-23 %) | *P*<0.01<br>-10 ± 3<br>(-20 %) | *P*<0.0001<br>-26 ± 6<br>(-25 %) | *P*<0.01<br>-57.67 ± 18.94<br>(-37%) |
| CWWs | Maximal amp<br>*F/Fo* | Duration<br>*ms* | Trise<br>*ms* | Tdecay<br>*ms* |  |
| **Normal**<br>(7 hearts)<br>(mean ± SE) | 7.3 ± 0.8<br>(n=14 transients) | 410 ± 45<br>(n=14) | 128 ± 22<br>(n=14) | 282 ± 39<br>(n=14) | |
| **MI**<br>(6 hearts)<br>(mean ± SE) | 6.2 ± 0.5<br>(n=30 transients) | 317 ± 21*<br>(n=30) | 103 ± 10<br>(n=30) | 214 ± 14*<br>(n=30) | |
| **MI vs Normal**<br>*Difference*<br>(%Normal) | *ns*<br>— | *P* < 0.05<br>-92 ± 43<br>(-23%) | *ns*<br>— | *P*<0.05<br>-68 ± 33<br>(-24%) | |

### Numerical evidence of SR-Ca^2+^-uptake implication in the elevation of Ca^2+^ release

We postulated that the observed faster Ca^2+^ extrusion from the cytosol reflected an increased SR-Ca^2+^ uptake in the MI Pcells. Analysis of wavelet transients also revealed a significant increase of the Ca^2+^ transient amplitude (**Fig.1B**.a, Tab. 1), suggesting that alteration of SR-Ca^2+^ uptake was accompanied by an elevation of SR-Ca^2+^ release. This raised the question as to whether the increase of SR-Ca^2+^ release responsible for larger SCA in MI Pcells is a primary alteration of myocardial ischemic conditions or a secondary effect of the SR-Ca^2+^-uptake elevation. We addressed this question by using a numerical model previously developed to mimic the intracellular Ca^2+^ mobilization of Purkinje fibers ^18,27^. As shown in **Fig.2A**, the model accurately reproduced the time course of both normal and MI Ca^2+^ wavelets as it was captured by LSCM in this study (see also supplement **Fig.3S**). The mean of maximal rates of Ca^2+^ uptake (*Umax*) calculated for each simulation of individual wavelets (see supplement **Fig.3S**) revealed a 52% increase in simulated MI transients compared to simulated normal transients (shown in supplement **Fig.3S**.E). However, as indicated in **Fig.2B**, incremental increase of *Umax* (**Fig.2B**.a) in the numerical expression of averaged normal transient shown in Figs.2A.a and 2B.c (blue trace) indicated that 35% elevation of *Umax* (**Fig.2B**.a) was sufficient to reproduce the transient of MI wavelets (see red trace in **Fig.2B**.c). No alteration of other model input parameters (e.g., diffusion coefficient, Ca^2+^-release threshold, etc.; see supplement “model overview”) was required. Interestingly, each *Umax* increment predicted a simultaneous rise in the amplitude of Ca^2+^-release pulse (**Fig.2B**.b), thus supporting the idea that increased Ca^2+^-release is a consequence of the elevation of SR-Ca^2+^-uptake in MI Pcells.

**FIGURE 2.**
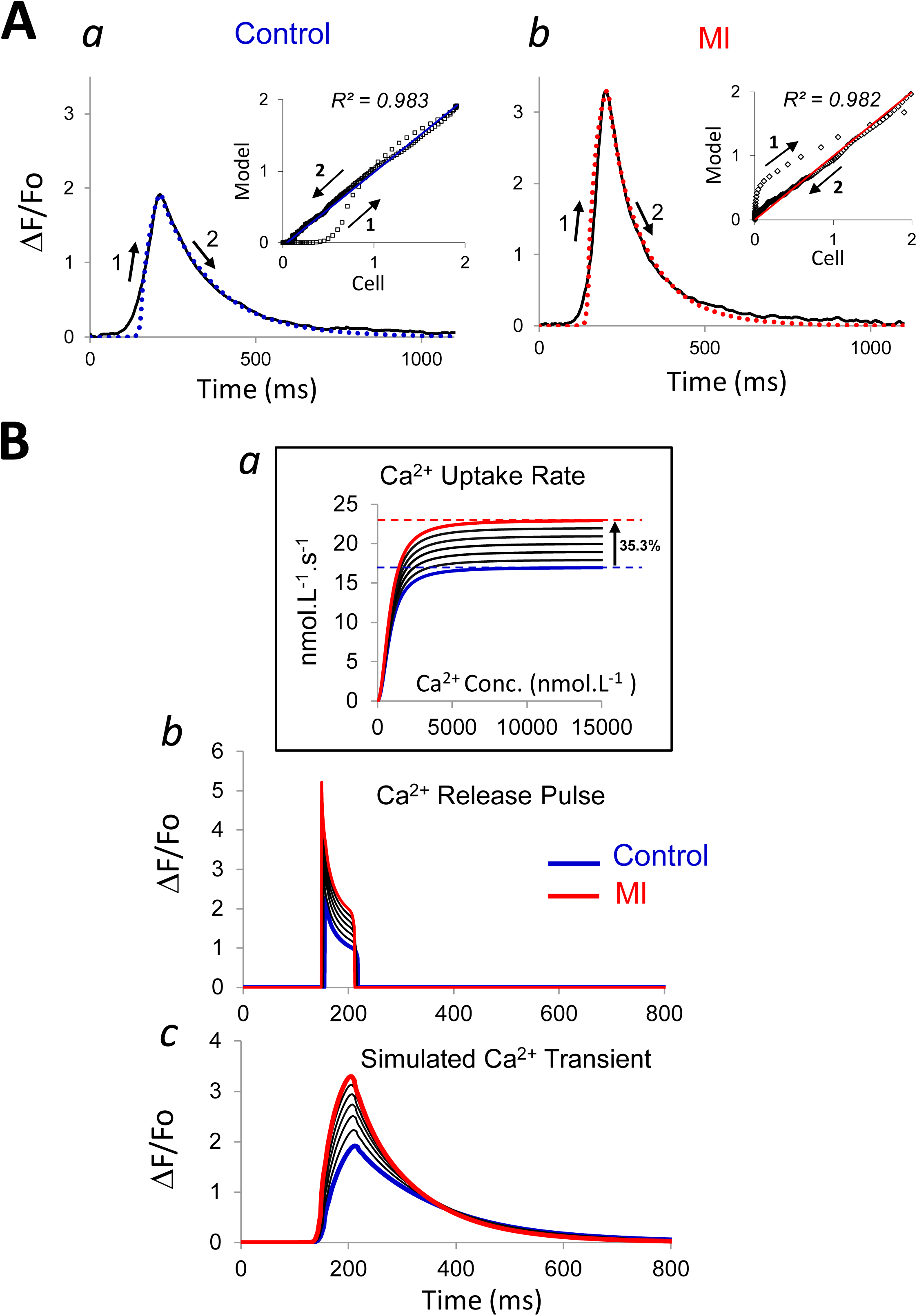
SR-Ca^2+^-uptake implication in Ca^2+^ transient alterations of MI Pcells. **A: Modeling of averaged wavelet Ca^2+^ transient:** 10 and 17 representative wavelets were combined, respectively, to produce the averaged Ca^2+^ transients of normal (**panel a**) and MI (**panel b**) Pcells; a Purkinje-specific model of intracellular Ca^2+^ mobilization (see text and “overview of the model” in supplement) was used to replicate the averaged transient in panels a and b; Numerical (dotted line) and experimental (plain line) curves were superimposed and accuracy of the simulations was estimated by linear regression as indicated in insets; rising and decay phases of the transients are indicated on the curves by 1 and 2, respectively. **B: Prediction of the effects of increase of SR-Ca^2+^-release on Ca^2+^ transient of normal cells**. Rate of Ca^2+^-uptake was progressively increased in the range of Ca^2+^ concentrations measured during the Ca^2+^ transient by incremental elevation of Vmax (**panel a**); the simultaneous effects on Ca^2+^ release pulse and overall transient were represented in panel b and panel c, respectively; in panel c, complete switch from normal transient (blue trace) to MI transient (red trace) was achieved with ∼35% increase of Vmax (panel a). The model description is detailed in the supplement.

### Decrease of SERCA pump density in Pcells after MI

Ca^2+^ transient analyses strongly suggest an alteration of SR-Ca^2+^-uptake function in Pcells after MI. Therefore, we examined the integrity of the SR-Ca^2+^ uptake system in canine MI Pcells by mapping the density distribution of SERCA2 by immuno-fluo-detection. It is known that phosphorylation of the regulatory protein phospholamban (PLB) increases the affinity of the SR-Ca^2+^ ATPase for Ca^2+^ and activates the pump function of the molecule ^31^. For this reason, SERCA2 was co-stained with pPLBSer16, phosphorylated form of PLB (pPLB). The distribution of SERCA2 antibody in Pcells of normal ventricle was transversally uniform (**Fig.3B**.a) and exhibited the typical sarcomeric striation previously described ^27^ (**Fig.3A**.a, **Fig.4A** (left panel)). The pPLBSer16 staining showed similar uniform transverse arrangement (**Fig.3A**.c, **Fig.3B**.c) and co-localized with SERCA2 antibody throughout the cell (**Fig.3A**.e). In contrast to normal Pcells, in 70-80% of MI Pcells (from infarcted ventricles) the SERCA2 staining intensity showed a large negative gradient from peripheral to central regions (**Fig.3A**.b, **Fig.3B**.b, **Fig.4A** (right panel); see also supplement Figs.4S). In these MI Pcells, pPLBSer16 staining exhibited the same intensity and uniform distribution compared to that seen in normal cells (**Fig.3A**.d, **Fig.3B**.d). SERCA2 and pPLBSer16 co-localized exclusively at the cell periphery (**Fig.3A**.f).

**FIGURE 3.**
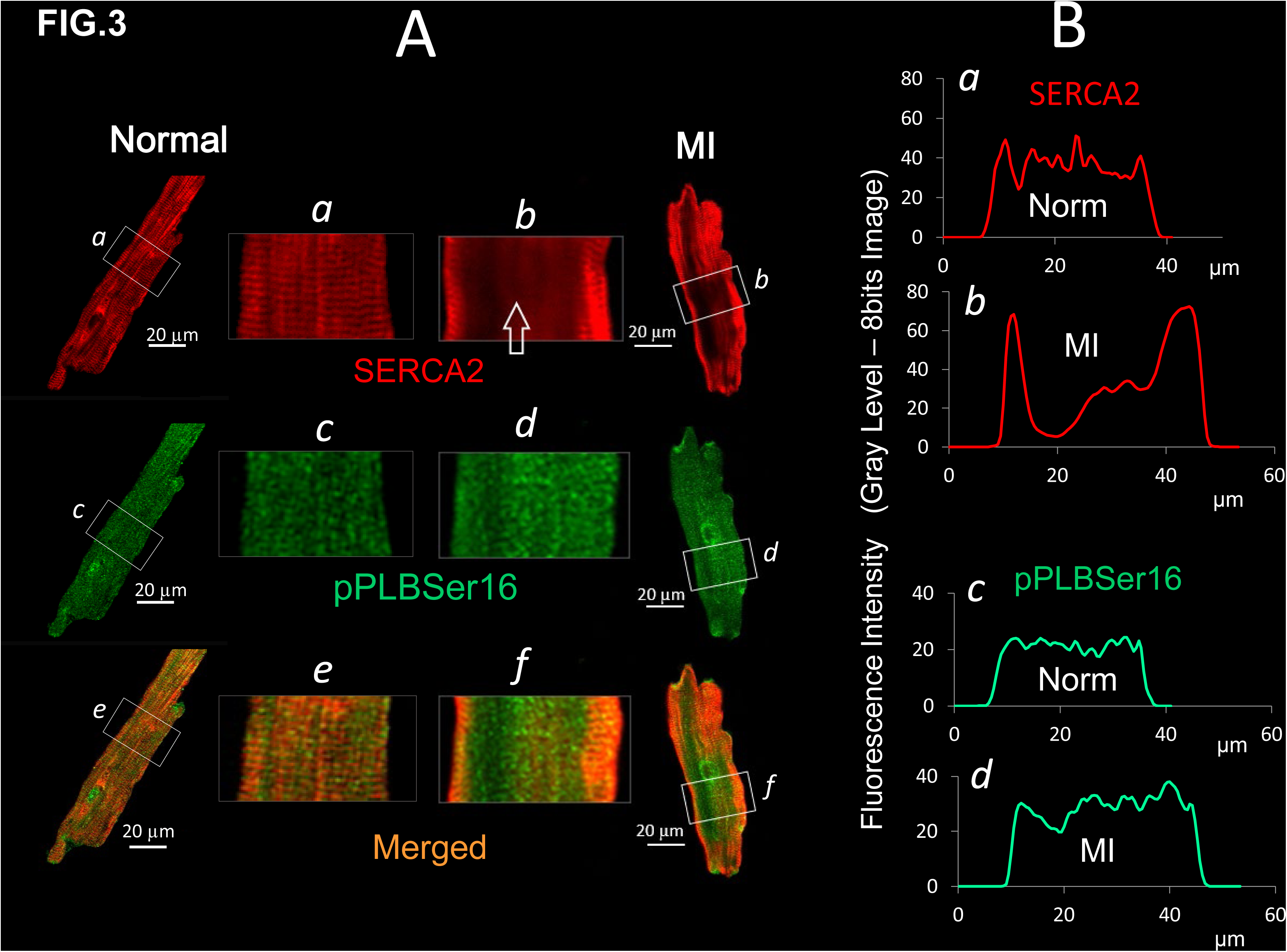
Immuno-characterization of SR-Ca^2+^ uptake in Pcells of dog heart with MI. **A. Typical immunostaining of SERCA2 and pPLBSer16**: SERCA2 pump and phosphorylated PLB (pPLBSer16) were co-stained in single Pcells of dog normal heart (*a and c*) and dog heart with MI (*b and d*); the degree of colocalization of SERCA2 and pPLBSer16 is shown in *e* and *f*, respectively. **B. Transverse distributions of SERCA2 and pPLBSer16 antibodies**: Distribution of antibodies shown in **A** was measured in normal (a and c) and MI (b and d) Pcells by averaging the 15 horizontal pixel profiles of areas *a, b, c*, and *d* selected in **A**, respectively. Note the staining deficit of SERCA2 in *b* compared to *a* while pPLBSer16 distribution in *d* is unchanged compared with *c*.

**FIGURE 4.**
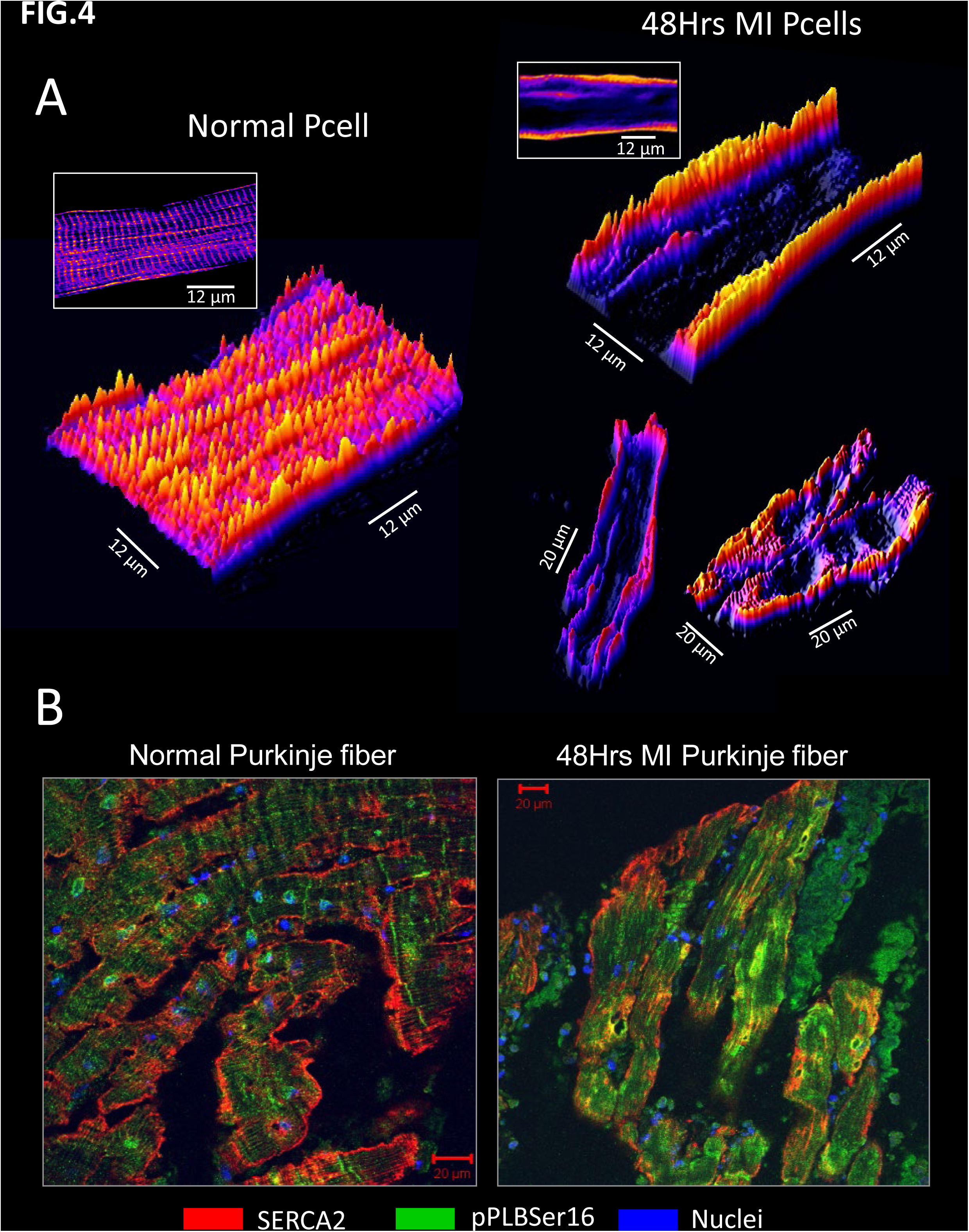
Expression of SERCA2 distribution heterogeneity in MI Pcells. **A. Two-dimensional representations of the two different distributions of SERCA2 density:** typical example of uniform sarcomeric arrangement of SERCA2 in normal Pcells (left) is compared with representative examples of heterogeneous distribution in MI Pcells (right); Note the large staining deficit in the core compared to the edges of MI Pcells. **B. Deficit of SERCA2 compared to pPLB in the Purkinje tissue of dog MI heart**: the ratio pPLBSer16/SERCA2 is two times larger in MI Purkinje tissue (right panel) compared to normal (left panel) due to decrease of SERCA2 staining in the core of MI Pcells (see green dominant pPLB staining in right panel): 0.058 ± 0.015 in normal Purkinje fibers (n=4 slices, N=3 hearts) versus 0.119 ± 0.019 in MI Purkinje fibers (n=6; N=3; P<0.05, powered 0.44).

Pixel analysis (see methods) of Purkinje tissue treated with the same antibodies (**Fig.4B**) revealed a twofold increase of the ratio pPLBSer16/SERCA2 expression in Purkinje tissue post MI. When compared with **Fig.3**, this result is consistent with the decline of SERCA2 staining while pPLB expression remains unchanged in the core of MI Pcells.

To detect a potential specificity of the canine model in our observations, we carried out SERCA2 immuno-staining in single Pcells of sheep (**Fig.5**) and human (supplement **Fig.5S**) hearts with comparable ischemic ventricular infarction (see **Fig.5A** and supplement **Fig.5SA**). SERCA2 antibody exhibited a uniform transverse distribution in the majority of Pcells of sheep (**Fig.5B**.a) and human (supplement **Fig.5SB**.a) normal hearts. In contrast, a pronounced deficit of SERCA2 staining comparable to that evidenced in the canine model was observed in the core of most Pcells of sheep (**Fig.5B**.b) and human (supplement **Fig.5SB**.b) hearts with MI. In a separate group of sheep Pcells, SERCA2 was co-stained with an important element of the SR membrane, the ryanodine receptor RyR2 (see supplement **Fig.6S**). While identical decline of SERCA2 staining was present in the core of sheep MI Pcells, no modification was found in RyR2 staining compared to normal cells, thus excluding a decreased SR density or restricted access for antibodies from potential causes of lower SERCA2 staining in the core.

**FIGURE 5.**
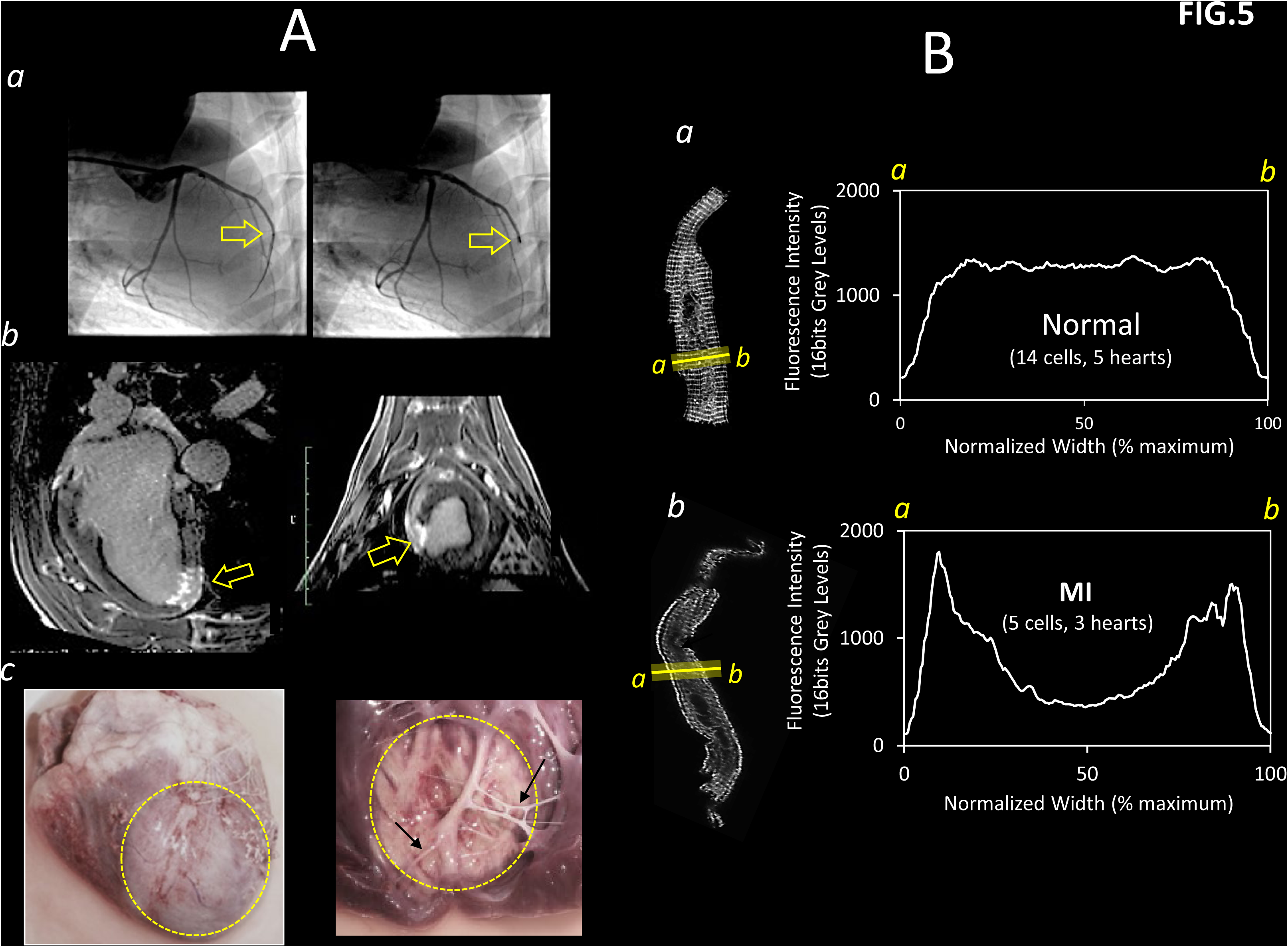
Effect of acute ischemic MI on SERCA2 expression in Pcells of sheep heart. **A. Myocardial infarction induced by angiographic method in sheep heart**: MI was induced by positioning an occluded stent in the LAD coronary artery through transfemoral catheterization and X-Ray imaging (**panel a**); resulting MI was assessed in living animal by MRI (**panel b**); typical external and internal view of transmural ischemic MI is given in **panel c** (yellow circle). **B. SERCA2 distribution in sheep Pcells**: Purkinje fibers (see arrows in **A**, panel c) were dissected and Pcells (see representative examples shown in B), were isolated 48Hrs after coronary occlusion; Pcells from normal heart (**panel a**) and from heart with MI (**panel b**) were stained with SERCA2 antibody (see text); pixel intensity was measured along segments *a-b*; transverse profiles of individual Pcells were normalized to maximal cell width and combined to generate the typical (averaged) SERCA2 distribution of normal (**panel a**) and MI (**panel b**) Pcells.

### Quantitative and cell population analyses of SERCA2 protein distribution

To estimate the magnitude of SERCA2 protein re-arrangement in Pcells after MI, we compared the SERCA2 staining intensity at the periphery with that in the center of sheep and human cells (**Fig.6**). We used the ratio R = F_P_ /F_C_ where Fp and Fc were the fluorescence intensities measured, respectively, in peripheral and central regions of the same cells (**Fig.6A**.a). R showed, on average, a fourfold and sixfold increase in sheep and human MI Pcells, respectively (**Fig.6A**.b). Three distinct types of SERCA2 transverse distribution were identified in the whole population of Pcells (**Fig.6B**.a): Type1 showed no difference in SERCA2 staining intensity between the edge and the center (R<1.5); Type3 was characterized by a pronounced drop from the edge to the core (2.5<R); Type2 was intermediate between Type1 and Type3, showing a moderate decrease in the core relative to the edge (1.5<R<2.5). As shown in **Fig.6B**.b, Type3 staining was mostly found in Pcells of sheep and human infarcted ventricles but also in 7% of sheep and 17% of human normal Pcells. Type1 and, to a lesser extent, Type2 patterns were detected only in normal Pcells. Our approach indicated that depression of SERCA2 staining was primarily a characteristic of MI Pcells population. The presence of Type2 and Type3 in few Pcells of normal ventricles is unclear but an effect of local hypoxic/ischemic conditions during heart, tissue and/or cells handling cannot not be completely excluded. This eventuality would demonstrate the extreme rapidity and sensitivity of SERCA2 re-arrangement in Pcells.

**FIGURE 6.**
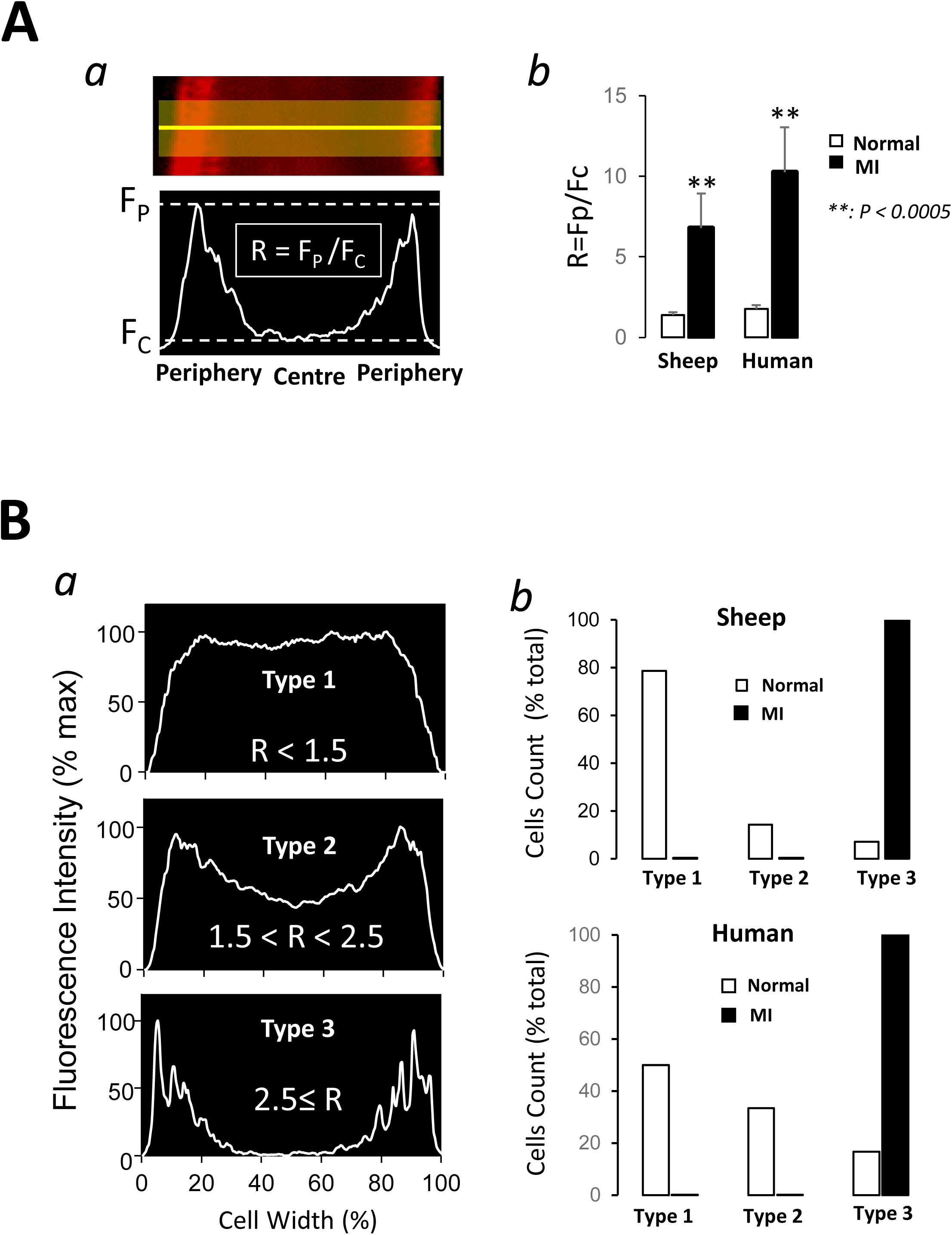
Quantification of SERCA2a distribution in sheep and human Pcells. A. **Quantification of the SERCA2 staining gradient**: The difference of SERCA2 staining intensity between peripheral and central regions of Pcells was measured in a transverse segment of the cell by R = Fp/Fc as indicated in **panel a;** Fp and Fc: fluorescence intensities measured, respectively, in the periphery and in the center of the same cell; large increases was detected in R value in both sheep and human Pcells after MI (**panel b**). B. **3 types of SERCA2 distribution:** As indicated in **panel a**, based on R value, three different types of transverse SERCA2 distribution were identified in the whole population of Pcells; **panel b**: in both sheep and human hearts, Most Type1 was found in normal cells whereas Type3 was seen most exclusively in MI cells (see text for details).

### SERCA2 gene/protein expression in LV infarcted myocardium

The apparent paradox between increased SR-Ca^2+^-uptake and decreased SERCA2 protein levels in Pcells of infarcted hearts prompted us to consider a potential compensatory modification of the pump isoform expression in ischemic ventricle (see discussion). We examined this hypothesis in adult Yucatan pig, a well-known model of human cardiac physio-anatomy. As shown in **Fig.7A**, ligation of the LAD coronary artery induced a 48Hrs infarct with location and anatomy comparable to other models. Tissues of infarcted LV exhibited a fourfold reduction of the SERCA2 transcript level compared to tissues of normal LV (**Fig.7B**.a). This result was consistent with the decrease of SERCA2 pump density detected by immunofluorescence in Pcells of canine infarcted heart. Analyses of SERCA2 sub-isoforms expression by PCR in the same tissues revealed a 85% drop of SERCA2a but also a 150% increase of SERCA2b transcripts compared to normal LV (**Fig.7B**.a). Matching the variations of transcript levels, immunoblots of LV tissues showed 59% decrease of SERCA2a protein level and a dramatic 967% increase of SERCA2b protein level in the infarcted LV compared to non-infarcted LV (**Fig.7B**.b),

**FIGURE 7.**
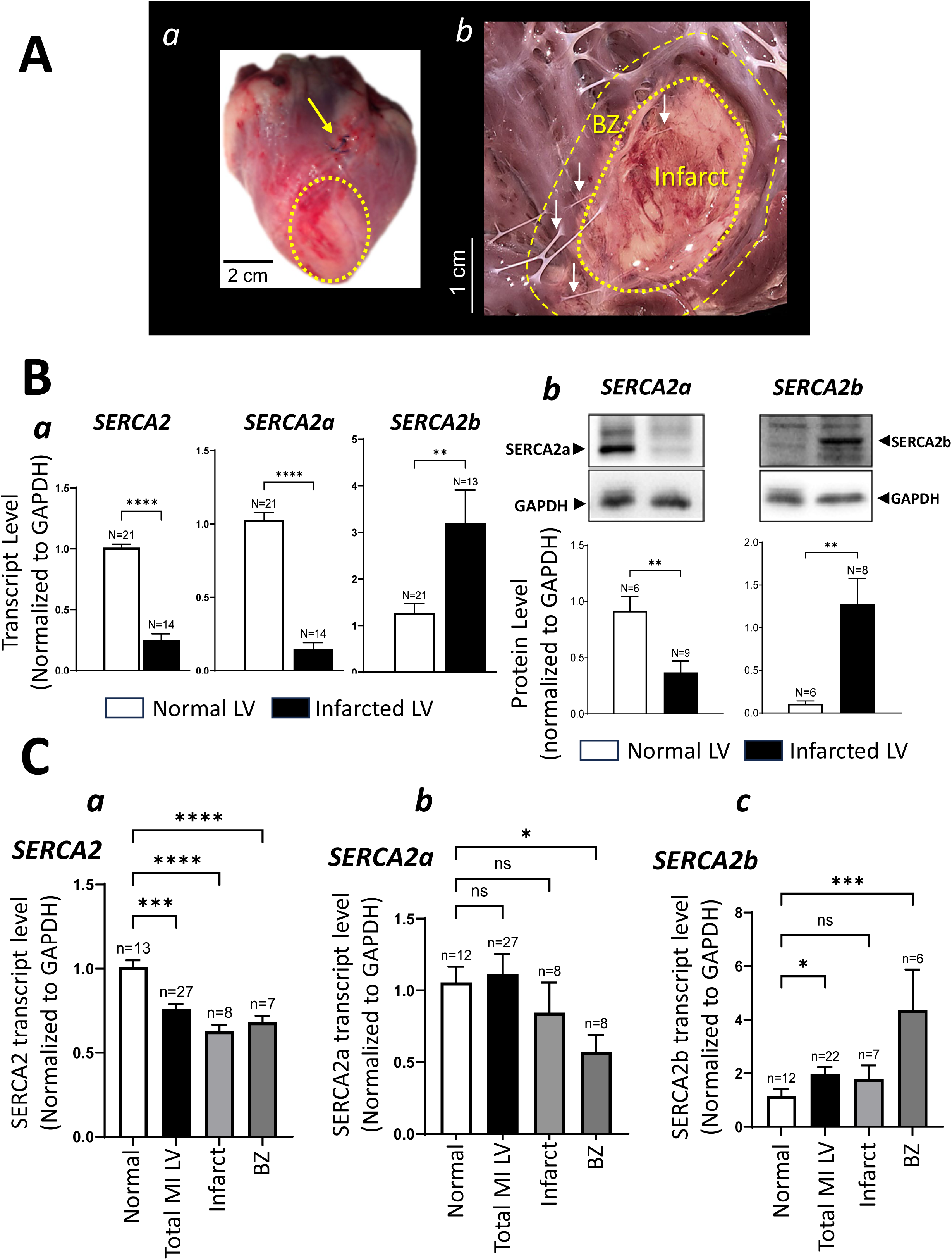
Analyses of gene/protein expression of SERCA isoforms in pig non-infarcted and infarcted ventricles. **A. Ischemic MI in pig heart:** MI was induced in pig heart by permanent ligation of the LAD artery (see arrow in **panel a**) following the same technique compared to dog model, leading to comparable transmural MI in the LV (see yellow circle in **panel a**); Purkinje fibers were dissected from the subendocardial region of the whole LV chamber; results from Purkinje fibers dissected in specific sub-regions referred as “Infarct” and “BZ” (see arrows in **panel b**) were considered separately. **B. Overall SERCA2 and SERCA2a/b expression in LV tissue**: transcript level was measured by RT-qPCR for SERCA2 and sub-isoforms 2a and 2b in tissues sampled from non-infarcted and infarcted LV (**panel a**); to evaluate the impact on protein expression of changes in splicing of SERCA2 pre-mRNA shown in **panel a**, SERCA2a and SERCA2b protein levels were measured by WB and reported in **panel b**. **C. SERCA2 and SERCA2a/b transcripts in Purkinje fibers:** SERCA2 and sub-isoforms transcripts levels were measured by RT-qPCR in Purkinje fibers isolated from normal and infarcted ventricles; in MI LV, the transcript level was evaluated in the total Purkinje fibers collected in the ventricle and in Purkinje fibers specifically located in the infarct and BZ regions. *: P<0.05 **: P<0.01; ***: P<0.001; ****: P<0.0001 ; ns: non-significant difference.

### SERCA2 gene expression in Purkinje fibers of infarcted LV

Purkinje strands were harvested from ventricles of 11 normal pig hearts and 12 hearts with 48Hrs MI. The levels of SERCA2 and SERCA2a/b transcripts in Purkinje samples were measured by RT-PCR and compared between infarcted LV and normal LV. In the pool of fibers collected from infarcted LV, subsets of fibers dissected specifically from the ischemic region (infarct) and border zone (BZ) (see **Fig.7A**.b) were examined separately. Consistent with the drop of overall SERCA2 gene expression in tissues of the infarcted LV (**Fig.7B**.a), we found a 24% reduction of SERCA2 transcript in the population of Purkinje fibers isolated from the infarcted ventricles (**Fig.7C**.a). This reduction was even more pronounced in Purkinje fibers of the infarcted region (37% decrease in the infarct and 32% in BZ). SERCA2a mRNA showed no variation in the total pool of fibers isolated from the infarcted LV (**Fig.7C**.b) whereas a 70% augmentation was detected in SERCA2b mRNA (**Fig.7C**.c). Although Purkinje fibers from the infarcted area exhibited a reduction of SERCA2a and an augmentation of SERCA2b transcripts, these variations were not statistically significant. However, the fibers spanning specifically the BZ exhibited a significant 46% decrease in SERCA2a transcript and a dramatic 280% increase of SERCA2b transcript.

## Discussion

### Increase of SR-Ca^2+^ uptake in Pcells of infarcted ventricles

We found that SR-Ca^2+^ release channels fired Ca^2+^ sparks at 60% higher rate in Pcells of canine ischemic heart compared with those of normal heart. This result is consistent with the elevation of pro-arrhythmic spontaneous Ca^2+^ activity (SCA) reported before in Purkinje fibers of ischemic heart ^19,20^. An increase of Ca^2+^ sensitivity of SR-Ca^2+^ channels (e.g. due to lower luminal Ca^2+^ activation threshold ^33^) was proposed to explain the elevation of SCA in canine MI Pcells ^20^. However, we report here that in the same canine MI cells the density of active Ca^2+^ release sites was unchanged whereas larger channel Ca^2+^ sensitivity should promote the activation of a larger number of Ca^2+^ release channels. In fact, the increased Ca^2+^ sensitivity of SR-Ca^2+^-channels was initially proposed for the interpretation of an upward shift of the curve Ca^2+^ event frequency versus Ca^2+^ concentration (pCa) in MI Pcells ^20^. But, the shifted relationship was detected in both peripheral and central regions of Pcells. It is unlikely that, within 48Hrs, the same alteration affects simultaneously the Ca^2+^ release threshold of the different Ca^2+^ release channels (RyR2, RyR3) previously reported in those subcellular regions of Pcell ^18,22^. Alternatively and consistent with an upward shift of Ca^2+^ event rate-pCa relationship, an acceleration of Ca^2+^ recycling by the SR can increase the opening frequency of a constant number of SR-Ca^2+^ channels. Free-Ca^2+^ circulates faster in the SR in two situations: (1) reduction of Ca^2+^ buffering by the reticular proteins such as CASQ2 ^34^ and (2) augmentation of SR-Ca^2+^-uptake by the SERCA pumps ^35^. Identical caffein-induced Ca^2+^ transient was found in normal and MI canine Pcells ^20^, indicating that the amount of mobilizable Ca^2+^ in the SR compartment is unchanged post MI, thus ruling out an alteration of reticular Ca^2+^ buffering function. However, a “stronger” Ca^2+^ sequestration by the SR is a plausible cause for faster SR-Ca^2+^ recycling. We found a 37% acceleration of Ca^2+^ decline during the wavelet transient, confirming the previous observation of “micro-Ca^2+^ transients” in canine MI Pcells by Hirose and colleagues ^20^. The decay phase of Ca^2+^ transients directly reflects the Ca^2+^ extrusion from the cytosol. In our study, the same decay shortening was seen in transients generated in central (CWWs) and peripheral (wavelets) regions, indicating that an identical and ubiquitously distributed Ca^2+^ transport system was responsible for the acceleration of cytosolic Ca^2+^ extrusion. In normal Pcells, the absence of T-tubules ^22,23,36,37^ implies that Ca^2+^ sequestration by the SR dominates the cytosolic Ca^2+^ extrusion in the core. Consequently, an increase of Ca^2+^-uptake by the SR-Ca^2+^-ATPases is a suitable interpretation of the acceleration of transient decay evidenced in wavelets and CWWs of MI Pcells.

### Increased SR-Ca^2+^ uptake and Ca^2+^ arrhythmogenicity in Pcells post MI

A computational model of intracellular Ca^2+^ mobilization in Purkinje fibers ^18,27^ enabled us to verify whether increase of SR-Ca^2+^-release in the cell periphery as suggested by the elevation of wavelet amplitude (Tab. 1) could directly result from the increased SR-Ca^2+^-uptake. This test successfully supported the idea that increased SR-Ca^2+^-uptake is a primary alteration associated with ischemia and could be the actual source of elevated SCA in MI Pcells.

We found that same alteration of SR-Ca^2+^-uptake at the periphery and in the core of MI Pcells, apparently affected the amplitude of the peripheral wavelets but not the central CWWs. As shown by immunostaining, a unique and ubiquitously distributed SR-Ca^2+^ ATPase (SERCA2) controls the SR-Ca^2+^-uptake throughout Pcell. However, SR-Ca^2+^-release is regionally controlled by 2 different channels: RyR2 in the core, and RyR3 at the periphery ^18,22^. In situ, RyR3-Ca^2+^-release showed evidence of larger Ca^2+^ sensitivity compared to RyR2 ^18,22,27^. Under these conditions and as it was developed in ^22^, identical increase of Ca^2+^-uptake from a continuous SR compartment is predicted to activate more Ca^2+^ release at the periphery than in the core. This prediction matches here our analysis of regional Ca^2+^ transients. This regional distinction was not considered in spark analysis of Fig.1 since line-scans were positioned randomly throughout the cells and, therefore, reflected only the mean tendency of Ca^2+^ release sites activity in Pcells.

Our conclusions raise the question of the mechanism whereby increased SR-Ca^2+^-uptake promotes Ca^2+^ arrhythmogenicity in Pcells. As mentioned above, the apparent lower Ca^2+^-release threshold of RyR3 versus RyR2 ^18,22^, predicts that SR-Ca^2+^-uptake primarily augments the frequency of RyR3-Ca^2+^ release at the periphery and potentiate the incidence and amplitude of lateral wavelets. In turn, larger wavelet amplitude and incidence increase the RyR2-SR-Ca^2+^ release by “Ca^2+^-induce Ca^2+^-release” in the core, and augment the probability of electrogenic/arrhythmogenic CWWs ^18,22^.

### Decrease of SERCA2 protein density is a characteristic of Pcells of infarcted ventricle

Our study then addressed the origin of SR-Ca^2+^-uptake elevation in MI Pcells. Immunostaining revealed a dramatic reduction of SERCA2 protein expression in the core of canine, ovine, and human Pcells after MI. The unchanged density distribution of pPLB in Pcells of canine MI heart indicates that SERCA2 reduction was the most likely cause of the twofold augmentation of the pPLB/SERCA2 ratio measured in MI Purkinje tissues (**Fig.4B**). Also consistent with the decreased SERCA2 protein expression evidenced in Pcells, the level of SERCA2 transcript exhibited a significant drop in tissues of porcine infarcted LV. The fact that SERCA2 decreases whereas typical proteins of the SR membrane such as PLB (canine Pcells) and RyR2 (ovine Pcells) remained unaltered confirmed that decline of central SERCA2 protein staining was not due to lower SR density or limited antibody accessibility. Cell population analysis of sheep and human Pcells showed that central drop of SERCA2 expression was a common characteristic of Pcells isolated from infarcted ventricles. Nevertheless, there was an apparent contradiction in those cells between drop of SERCA2 density and elevation of SR-Ca^2+^-uptake. We propose that such paradox may reveal the existence of an intermediate feature between decreased SERCA pump density and increased SR-Ca^2+^ uptake. We hypothesized that this intermediate feature resides in the nature of SERCA2 sub-isoforms.

### Alteration of SERCA2 mRNA splicing in tissues of infarcted ventricle

In the heart, SR-Ca^2+^ ATPase is encoded by the SERCA2 gene (*ATP2A2*) ^38^. Tissue-specific alternative processing of SERCA2 pre-mRNA produces the SERCA2a protein in the heart, slow-twitch skeletal muscles, and smooth muscles, and SERCA2b virtually in all other tissues ^39^. Although human-specific splicing variants SERCA2c and SERCA2d have been reported ^38^ and SERCA2b have been found at low levels in the heart ^40^, SERCA2a represents more than 95% of SERCA pump isoforms in mammalian cardiac cells ^38,41^. Total or partial inhibition of cardiac SERCA2a expression in transgenic mice models *SERCA2^-/-^* and *SERCA2^+/-^* resulted in a total or partial increase of SERCA2b isoform and decrease of SERCA2 mRNA/protein expression in the heart ^42^. Same variations of SERCA2b and SERCA2 associated with the drop of SERCA2a expression were observed in our study (**Fig.7B**), thus suggesting that upregulation of SERCA2b and decrease of SERCA2 gene/protein expression in Pcells and tissue of infarcted LV, result from the downregulation of SERCA2a.

Expressed in COS cell system, SERCA2b was reported to have twice larger affinity for Ca^2+^ but half turnover rate for Ca^2+^ compared to SERCA2a ^43–45^. Nevertheless, in the same studies, further examination of Ca^2+^ uptake rate at a given pCa clearly suggests 2-3 times increase of Ca^2+^ transport rate for SERCA2b compared to SERCA2a in the range of Ca^2+^ concentrations (0.1-1μM) measured during the decay of Ca^2+^ transients (see Fig.4 in ^43^ and Fig.3 in ^44^). In addition, it was found in mouse working heart preparation that overexpression of SERCA2b in cardiac tissue was accompanied by a significant increase of ventricular relaxation rate (-dP/dt) ^46^. This increase is consistent with an augmentation of cytosolic Ca^2+^ removal in cardiac cells. Finally, considering that SERCA2b can functionally replace SERCA2a when it is targeted to myocardial SR ^42^, we propose that increase of SR-Ca^2+^-uptake rate in Purkinje fibers of infarcted heart could be the result of a switch in SERCA2 splicing leading to SERCA2a downregulation and SERCA2b upregulation. Assuming a larger Ca^2+^ transport rate of SERCA2b in vivo compared to SERCA2a and considering the dramatic augmentations detected in SERCA2b expression in our study, the upregulation of SERCA2b could be sufficient to compensate for the SERCA2 decrease. Equivalent conclusion was numerically predicted in ^47^. A potentiation of SERCA2b Ca^2+^ affinity by phosphorylated PLB ^44^ may also contribute to increased SR-Ca^2+^-uptake. This scenario is supported in our study by the larger pPLB/SERCA protein ratio in Purkinje fibers of infarcted LV, possibly highlighting an increased proportion of Ca^2+^ pumps activated by PLB phosphorylation.

## Conclusion

Our study supports the hypothesis that ischemia is accompanied by a change in the SERCA2 splicing process leading to an upregulation of SERCA2b and downregulation of SERCA2a. The measures of protein and gene expression support this hypothesis not only in the Purkinje fibers but also in the whole myocardium of the infarcted LV. The interesting finding that largest increase of SERCA2b expression was measured in Purkinje fibers spanning the border zone (BZ) is, potentially, a novel insight into the origin of specific arrhythmogenicity reported in this region ^48–51^.

By “boosting” the contractile properties of ventricular myocytes, the SERCA2b expression may reflect an immediate adaptation of cardiac muscle to the loss of regional contractility due to MI. This extremely rapid modification might be tissue non-discriminatory, affecting all cells in the ventricle including those of Purkinje fibers. Purkinje arrhythmogenicity may be seen as a negative side effect of this molecular adaptation of the heart to MI.

## Acknowledgments

We acknowledge the participation and support of animal care services of Memorial University, University of Calgary, and Liryc institute for all procedures involving animals. We acknowledge the surgery team (groupe hospitalier Pellegrin, Bordeaux) for the provision of human hearts.

